# The C-terminal extension landscape of naturally presented HLA-I ligands

**DOI:** 10.1101/213264

**Authors:** Philippe Guillaume, Sarah Picaud, Petra Baumgaertner, Nicole Montandon, Julien Schmidt, Daniel E Speiser, George Coukos, Michal Bassani-Sternberg, Panagis Fillipakopoulos, David Gfeller

## Abstract

HLA-I molecules play a central role in antigen presentation. They typically bind 9- to 12-mer peptides and their canonical binding mode involves anchor residues at the second and last positions of their ligands. To investigate potential non-canonical binding modes we collected in-depth and accurate HLA peptidomics datasets covering 54 HLA-I alleles and developed novel algorithms to analyze these data. Our results reveal frequent (442 unique peptides) and statistically significant C-terminal extensions for at least eight alleles, including the common HLA-A03:01, HLA-A31:01 and HLA-A68:01. High resolution crystal structure of HLA-A68:01 with such a ligand uncovers structural changes taking place to accommodate C-terminal extensions and helps unraveling sequence and structural properties predictive of the presence of these extensions. Scanning viral proteomes with the new C-terminal extension motifs identifies many putative epitopes and we demonstrate direct recognition by human CD8+ T cells of a C-terminally extended epitope from cytomegalovirus.

## Introduction

Human Leukocyte Antigen class I (HLA-I) molecules play a major role in immune defense mechanisms by presenting to T cells peptides from the intracellular matrix (1). Peptides presented on HLA-I molecules originate mainly from proteasomal degradation of self or pathogen-derived proteins. These peptides are first translocated to the endoplasmic reticulum. There, they can load on HLA-I molecules provided their sequence is compatible with HLA-I binding motifs. Peptide-HLA-I complexes are then transported to the cell surface where they can elicit T cell recognition, primarily upon presentation of non-self peptides.

Most HLA-I alleles preferentially bind 9- to 12-mer peptides, although important differences have been observed between alleles in terms of the preferred length of their ligands (2–6). The majority of alleles accommodate peptides with anchor residues at the second and last positions. From a structural point of view, anchor residues point directly into the HLA-I peptide binding groove. Their importance for HLA-I peptide interactions is also reflected at the sequence level, where alignments of HLA-I ligands display clear specificity at the second and last positions for most alleles. 9-mer HLA-I ligands are characterized by a linear binding mode. For longer peptides, numerous crystal structures have shown the presence of a bulge at middle positions, protruding outside of the HLA-I binding site in order to accommodate the additional residues between the two anchor positions (e.g., ref. (7, 8)).

Over the years, anecdotal evidences of C-terminal extensions beyond the last anchor position have been observed among HLA-A02:01 ligands and the first crystal structure with such a ligand was published in 1994 (PDB: 2CLR) (9). More recently, analysis of HLA peptidomics data obtained by Mass Spectrometry (MS) from cell lines transfected with soluble HLA-A02:01 and infected with *Toxoplasma gondii* revealed several C-terminal extensions and showed that, for this allele, C-terminal extensions were mainly found among peptides coming from pathogens and may involve cross-presentation (10, 11). X-ray crystallography revealed distinct structural mechanisms in HLA-A02:01 to accommodate C-terminal extensions (10, 11). N-terminal extensions have also been recently observed in HLA-B57:01 (12) and HLA-B58:01 (13). However, N- or C-terminal extensions have not been much investigated in other HLA-I alleles based on unbiased HLA-I ligand datasets (see some predictions for HLA-C04:01 in (14)). As such, it remains unclear whether they occur frequently and, if yes, whether they can be recognized by CD8 T cells.

Here, we introduce a novel statistical approach to rigorously investigate N- and C-terminal extensions in large datasets of naturally presented HLA-I ligands obtained by in depth and accurate HLA peptidomics profiling of cell lines and tissue samples covering more than 50 HLA-I alleles. Our work reveals widespread C-terminal extensions for at least eight HLA-I molecules (HLA-A02:03, HLA-A02:07, HLA-A03:01, HLA-A31:01, HLA-A68:01, HLA-A68:02, HLA-B27:05 and HLA-B54:01), and we identify both sequence and structural features in HLA-I alleles predictive of the presence of C-terminal extensions. A new crystal structure of HLA-A68:01 in complex with such a ligand uncovers structural changes to accommodate the C-terminal extensions. Scanning viral proteomes with our new motifs describing C-terminal extensions further enabled us to demonstrate direct CD8 T cell recognition of an HLA-A03:01 restricted C-terminally extended epitope from cytomegalovirus (CMV).

## Results

### Unbiased investigation of C- and N-terminal extensions

To investigate the presence of C- and N-terminal extensions in HLA-I ligands, we collected recent pooled and mono-allelic HLA peptidomics datasets from seven studies covering 43 samples, 54 different HLA-I alleles and 109,953 unique peptides (3, 15–20) (see Materials and Methods and Table S1). These datasets were all generated with <1% false discovery rate and were not filtered with any predictor. We hypothesized that N- or C-terminal extensions among naturally presented endogenous HLA-I ligands may be determined by identifying peptides of ten or more amino acids that do not match motifs expected for ligands following the bulge model (Figure 1A). For pooled HLA peptidomics data, 9-mer HLA binding motifs were identified and annotated with our recent motif deconvolution algorithm (4, 20) (see example in Figure 1B and Materials and Methods). Importantly, these motifs were identified without relying on HLA-I ligand predictors and therefore represent a fully unbiased view of the 9-mer HLA peptidome. Focusing on the first three and the last two positions in the 9-mer motifs, we then built Position Weight Matrices (PWM) modeling bulges, N-terminal extensions or C-terminal extensions for each of the 9-mer motifs (Materials and Methods and Figure 1C). For 10-mers, bulges were modeled by incorporating five non-specific positions between the first three and the last two residues. N-terminal extensions were modeled by adding four non-specific positions in the middle and one non-specific position at the N-terminus. C-terminal extensions were modeled by adding four non-specific positions in the middle and one non-specific position at the C-terminus (Materials and Methods and Figure 1C). Alleles displaying anchor residues at middle positions (e.g., HLA-B08:01) were not considered as 10-mer ligands are much less frequent for them (4).

**Figure 1.**
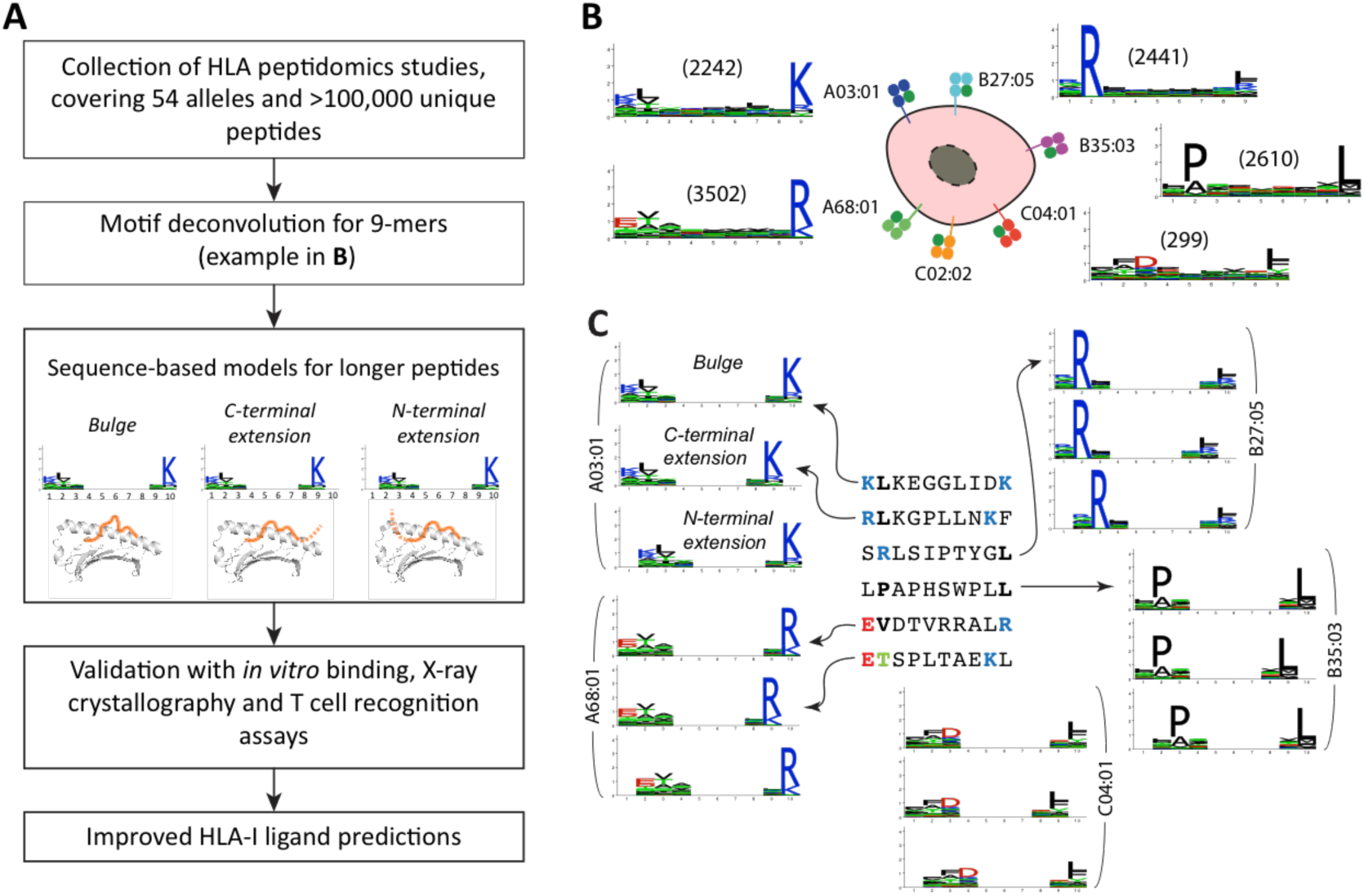
**A:** General description of the analysis pipeline developed in this work to identify and validate N- or C-terminal extensions. **B**: Example of 9-mer motifs identified in HLA peptidomics data from Mel_15 (15) with our motif deconvolution algorithm (4). The number of peptides assigned to each motif is shown with parentheses. **C**: Illustration of the different models built from the 9-mer motifs of panel B (bulge, C- and N-terminal extension) to investigate non-canonical binding modes among longer peptides (10-mers in this example).

Using the three models (bulge, N- and C-terminal extensions) derived from each motif observed among 9-mer ligands, we then scored all 10-mer peptides (see Figure 1C and Materials and Methods). Peptides that displayed a much higher score for exactly one allele and one model were assigned to this allele and the corresponding model. For the special case of mono-allelic cell lines (3), comparison was only performed among bulge, N- and C-terminal extension models of the same allele. To determine statistical significance of N- or C-terminal extensions, we further developed a null model representing the expected 10-mer HLA-I ligands assuming only bulges (Materials and Methods). We finally retrieved all sets of peptides predicted to follow N- or C-terminal extensions associated to a given allele in a given sample that could not be explained by statistical fluctuations in the data (Z-score > 2). For each set of peptides the motif was modeled with a PWM, graphically represented as a sequence logo. This resulted in 15 motifs of C-terminal extensions (for a total of 396 unique 10-mer peptides) and no motif of N-terminal extensions (Figure 2A and Table S3). Seven motifs corresponded to C-terminal extensions associated to HLA-A03:01 across different HLA peptidomics samples, two motifs to HLA-A68:01 and one motif to each of the other alleles (i.e., HLA-A02:03, HLA-A02:07, HLA-A31:01, HLA-A68:02, HLA-B27:05 and HLA-B54:01, Figure 2A, see also Figure S1, Table S2 and S3). We first note that motifs describing C-terminal extensions associated to the same allele (i.e., HLA-A03:01 or HLA-A68:01) in different datasets displayed a very high similarity among each other (rectangles in Figure 2A). This highlights the remarkable reproducibility of our predictions across our very heterogeneous set of studies performed in different places, with different protocols and on different cell lines or tissue samples. For all predicted C-terminal extensions that passed our statistical threshold, we show the average frequency of such ligands in Figure 2B (Materials and Methods). Overall, our results predicted widespread (i.e. from 3% to 16% of the 10-mer peptides) C-terminal extensions.

**Figure 2.**
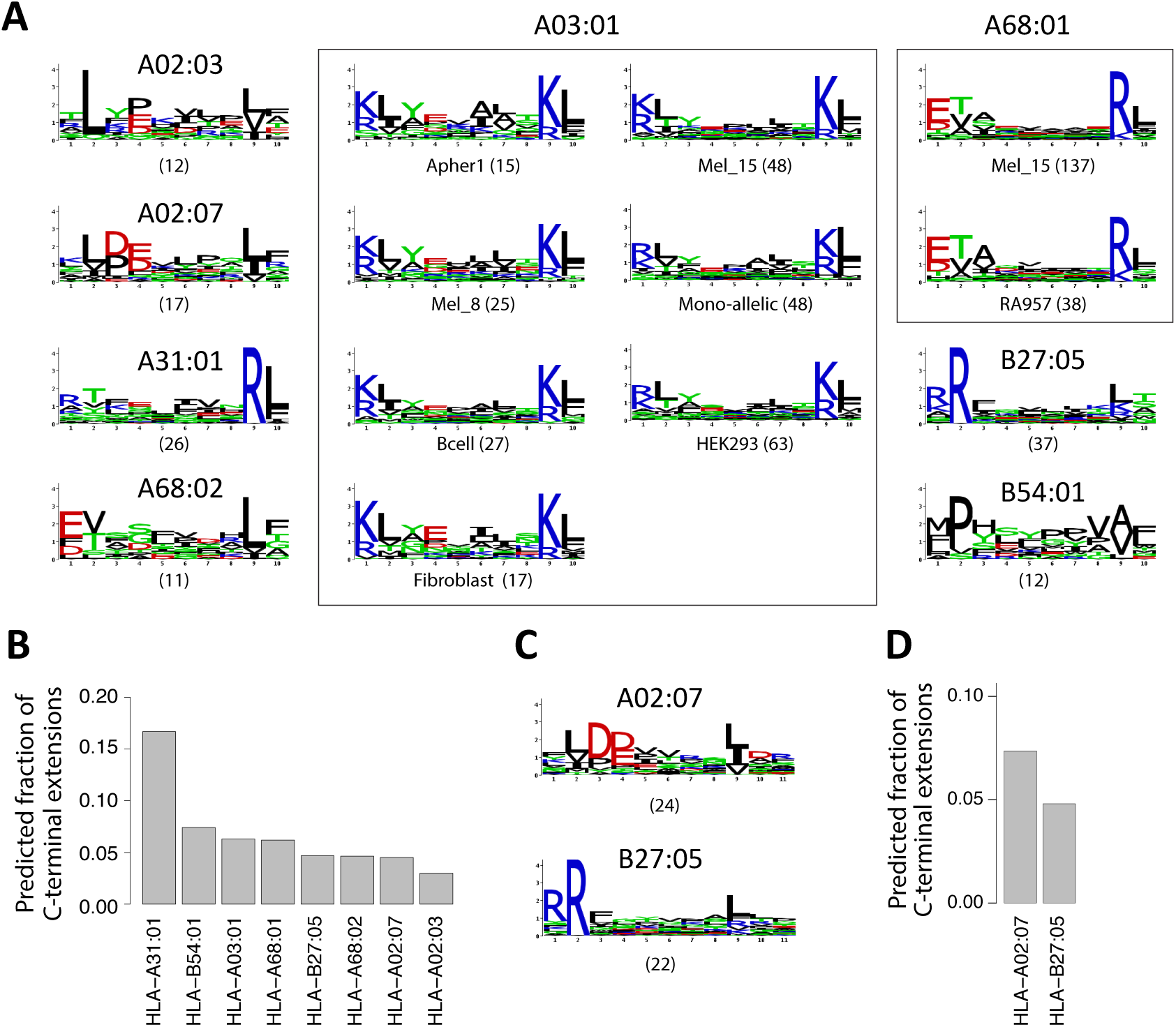
**A**: Predicted 10-mer C-terminal extension motifs found for different HLA-I alleles in the different datasets. Parentheses indicate the number of peptides predicted to follow the C-terminal extension model. Study of origin is indicated for motifs identified in multiple samples for the same HLA-I allele. **B:** Estimates of the frequency of C-terminal extensions among 10-mers for alleles shown in **A**. **C:** C-terminal extensions motifs comprising two residues after the second anchor residue in 11-mers ligands. **D:** Estimates of the frequency of C-terminal extensions among 11-mers for alleles shown in **C**.

To investigate whether C-terminal extensions may extend for more than one amino acid, we applied our approach to longer peptides (i.e., 11-mers, allowing for two unspecific positions at the N- or C-terminus) and found statistically significant evidences of C-terminal extensions for HLA-A02:07 and HLA-B27:05 for a total of 46 unique 11-mer peptides (Figure 2C-D, Table S2 and S3). Of note, the motifs are consistent with the 10-mer C-terminal extensions previously observed (Figure 2A). A trend was also observed for HLA-A02:01, HLA-A02:03, HLA-A03:01, HLA-A68:02 and HLA-B54:01 (see Table S3), although it did not pass our thresholds on the number of ligands or the Z-score, and was not consistently observed across samples. For other alleles, we did not observe statistical evidences of C-terminal extensions of two residues among 11-mers, although we cannot exclude that rare C-terminal extensions could be missed in MS data because of the competition with shorter ligands. When analyzing even longer ligands (12-mers), we did not find anything statistically significant, but the number of such ligands identified by MS is much smaller so that it may be difficult to confidently identify C-terminal extensions with our approach and distinguish them from potential noise in MS data.

### In vitro validation

To experimentally test our predictions, we selected several 10-mer ligands predicted to follow the C-terminal extension binding mode for three of the most frequent alleles with such predictions (i.e., HLA-A03:01, HLA-A31:01, HLA-A68:01). We then mutated either the last or second-to-last residue and experimentally measured the stability of the wild-type and the two mutated peptides (Materials and Methods). 9-mer peptides without the predicted C-terminal extensions were used as positive controls. As expected mutating the last residue of the 10-mer peptides had little effect on their binding affinity, while mutating the second to last residue significantly decreased the stability of the complexes (Figure 3). These data strongly suggest that, for all these 10-mer peptides, the second to last residue is playing the role of the anchor residue and the last residue is extending beyond the canonical C-terminus of HLA-I ligands. To test whether longer C-terminal extensions may bind to HLA-A03:01, we added all amino acids not compatible with the bulge model at the C-terminus of one of the C-terminally extended 10-mer ligands used in Figure 3 (KLAYTLLNKL*) and measured the stability of these 11-mer peptides (Figure S2). Our results indicate that most of the 11-mers did bind, although with lower stability compared to the 10-mer peptide (i.e. most complexes were fully dissociated at time 72h). This suggests that longer C-terminal extensions can extend for more than one residue in HLA-A03:01 ligands, although this likely corresponds to a small fraction of the actual HLA peptidome, as suggested by their low frequency in MS data.

**Figure 3.**
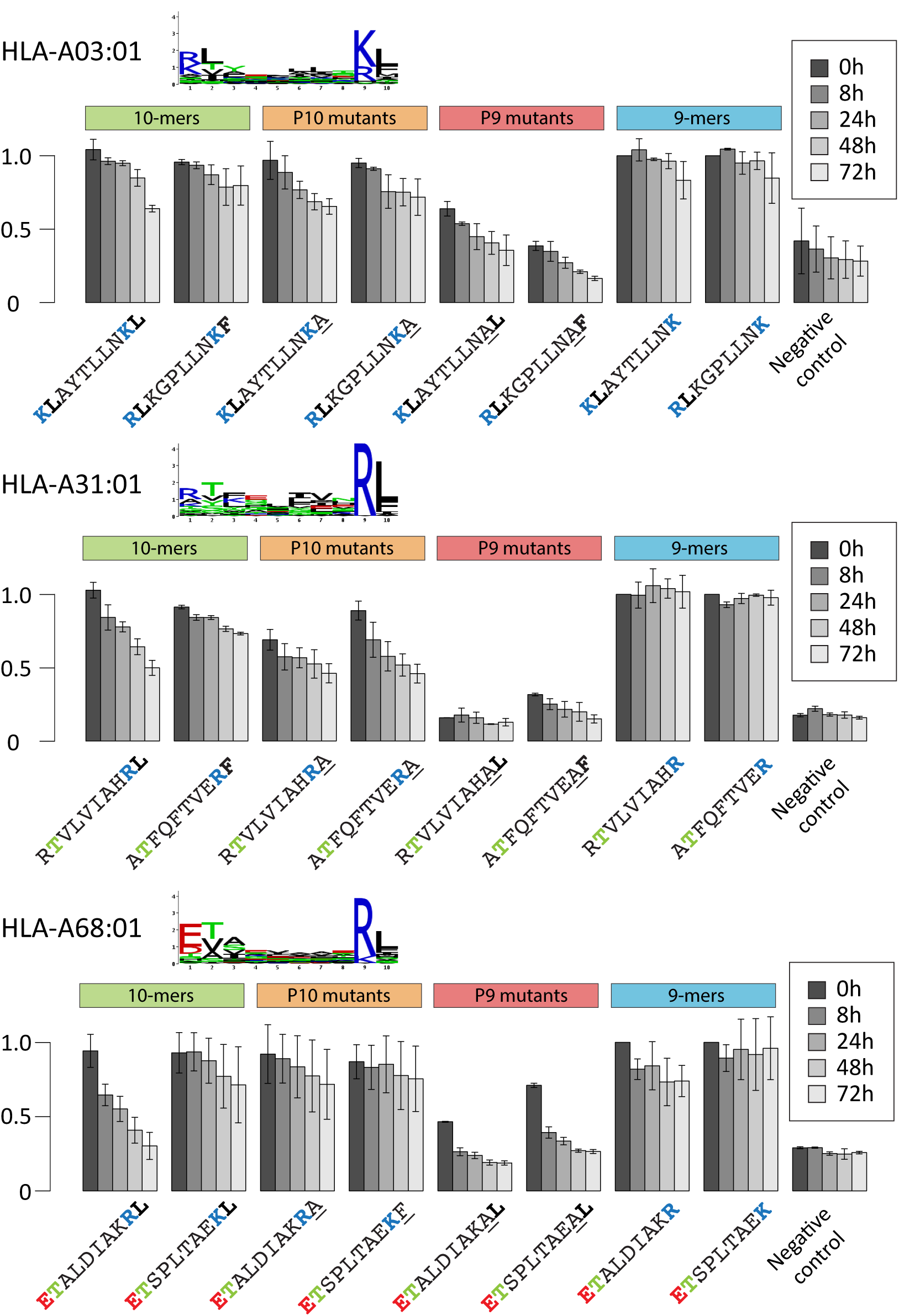
In vitro binding stability assays based on ELISA for peptides predicted to follow the C-terminal extension binding mode for HLA-A03:01, HLA-A31:01, HLA-A68:01 (mutated amino acids are underlined). The motif for the C-terminal extensions is shown for each allele.

### Robustness to noise

To explore the robustness of our findings with regard to noise in HLA peptidomics data, we rerun our whole pipeline (i.e., motif identification (4) and annotation (20) in 9-mers as well as C-terminal extension predictions in 10-mers) adding 5% of randomly selected peptides from the human proteome to all datasets. Remarkably, the predicted C-terminal extensions remained basically unchanged (Figure S3). In particular, we did not observe any new predicted C-terminal extension motifs that would arise by chance from the random peptides. This analysis clearly suggests that C-terminal extensions predicted in this work are not resulting from contaminations in HLA peptidomics data.

### Crystal structures of C-terminally extended HLA-I ligands

To investigate the structural mechanisms underlying C-terminal extensions uncovered in this work, we generated a high resolution (1.6Å) crystal structure of HLA-A68:01 in complex with a 10-mer peptide (`ETSPLTAEKL`, see Figure 3) predicted to follow the C-terminal extension binding mode (see Materials and Methods, Figure 4A and Figure S4). As expected, the lysine at P9 (yellow sidechain) filled the F pocket and superimposed nicely with the last residue (K9) of canonical 9-mer ligands of HLA-A68:01 (blue sidechain in Figure 4A, PDB:4HWZ) (21). More importantly, in order to accommodate the C-terminal extension (L10), the Y84 sidechain was flipped by 90 degrees and the two alpha helices around the F pocket moved away from each other (Figure 4A), as measured by the distance between C-alpha atoms of residues 80 and 143 (D_80-143_=11.1Å versus D_80-143_=10.1Å for HLA-A68:01 in complex with a 9-mer peptide). Interestingly, the same flip had been observed in one of the HLA-A02:01 structures in complex with C-terminally extended ligands (10) (see Figure 4B). However, in contrast to HLA-A02:01 where the sidechain of the first residue of the C-terminal extension points towards the solvent (K12 in PDB:5DDH in Figure 4B (10), see also L10 in PDB:5FA4 and D10 in PDB:5F7D (11)), in the case of HLA-A68:01, the C-terminal extension sidechain (L10) filled the pocket created by the flip of Y84 sidechain (see also Figure S5). This result is fully consistent with the specificity for hydrophobic residues in the C-terminal extension motif of HLA-A68:01 (Figure 2A).

**Figure 4.**
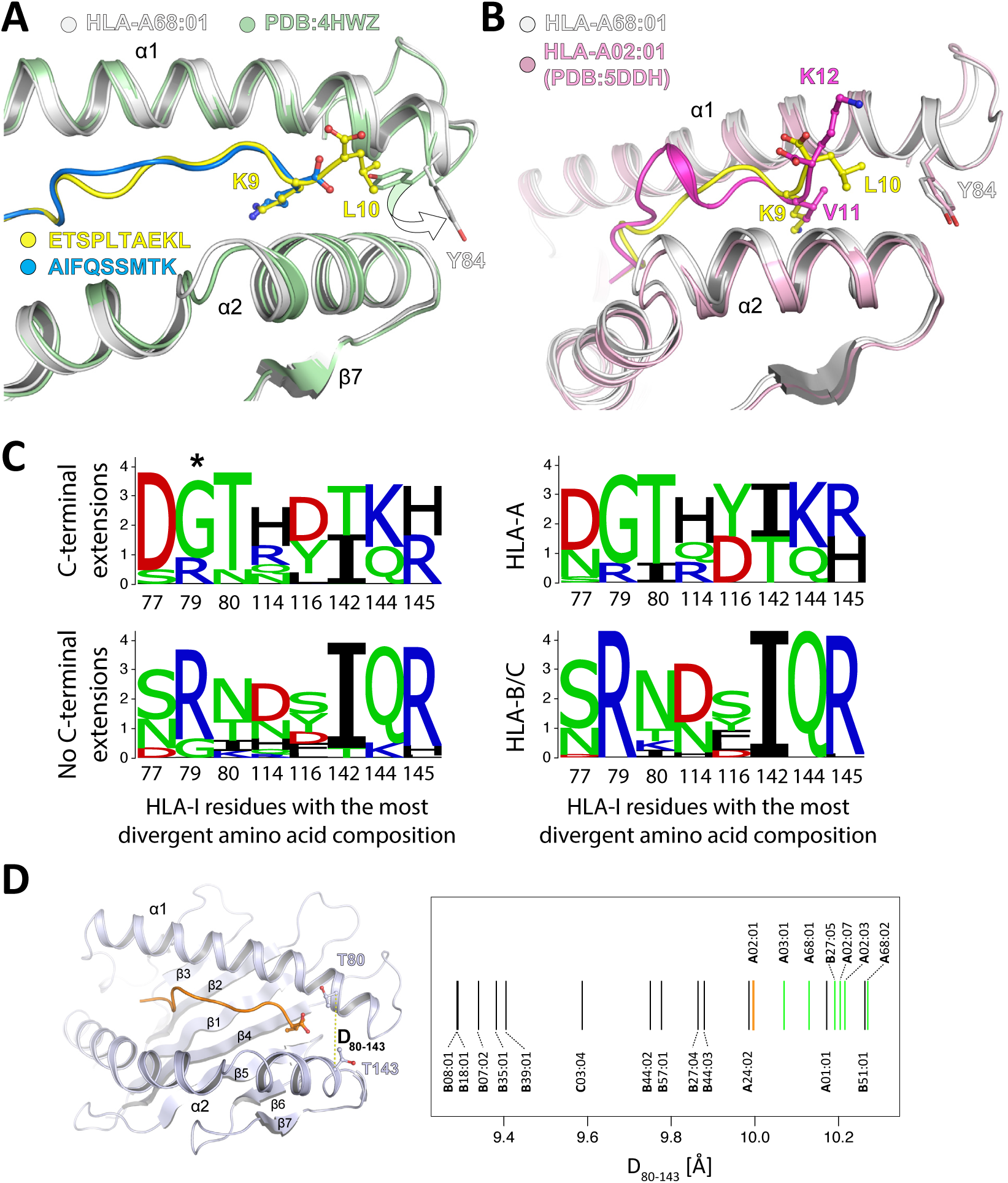
**A:** Crystal structure of HLA-A68:01 (white ribbon) in complex with a C-terminally extended ligand (yellow) superimposed with HLA-A68:01 (green ribbon) in complex with a canonical 9-mer ligand (blue, PDB:4WHZ) (21). **B:** Comparison of HLA-A68:01 (white ribbon) in complex with a C-terminally extended ligand (yellow) and the complex of HLA-A02:01 (pink ribbon) bound to a C-terminal extended 12-mer peptide (FVLELEPEWTVK, magenta, PDB:5DDH) (10). **C**: HLA-I residues surrounding the C-terminus of canonical ligands and displaying the largest Jensen-Shannon divergence between alleles with C-terminal extensions (top left) and alleles without C-terminal extensions (bottom left). For comparison, the sequence logos of HLA-A (top right) and HLA-B/C (bottom right) alleles at the same positions are displayed. **D:** Analysis of the distance between C-alpha atoms of residues 80 and 143 for alleles considered in this work and with available crystal structures (Table S4). Green lines correspond to alleles displaying C-terminal extensions. The orange line represents HLA-A02:01.

### Propensity of HLA-I alleles for C-terminal extensions

To investigate the molecular determinants of C-terminal extensions, we aligned the sequences of all HLA-I alleles considered in this work and checked whether some amino acid patterns in residues surrounding the F pocket would characterize alleles with C-terminal extensions (Materials and Methods). Clear differences were observed at specific positions (Figure 4C) and showed overlap with properties characterizing HLA-A alleles, as expected from the higher proportion of HLA-A alleles with C-terminal extensions. Interestingly, glycine is known to destabilize alpha helices and the presence of glycine at position 79 (star in Figure 4C) in most HLA-I alleles displaying C-terminal extensions may endow the α1 helix with the flexibility required to accommodate such extensions (Figure 4A). To test whether these amino acid patterns may help predict whether an allele is more likely to display C-terminal extensions, we trained a logistic regression and performed a rigorous cross-validation, using as features all amino acids surrounding the C-terminus of 9-mer ligands (Materials and Methods). An average Area Under the ROC Curve (AUC) of 0.85 could be obtained, suggesting that alleles accommodating C-terminal extensions may be reasonably well predicted from their sequence. Lower accuracy was reached in a simple model where HLA-A alleles are predicted to display C-terminal extensions and HLA-B/C alleles are not (AUC=0.76). To further investigate molecular mechanisms allowing for C-terminal extensions, we surveyed available X-ray structures of HLA-I alleles considered in this work (mainly in complex with canonical 9-mer ligands and not including any C-terminal extensions, see Table S4). We then computed the distance between the two alpha helices surrounding the C-terminus of canonical ligands (see before and Table S4). Interestingly, we observed that, on average, alleles with C-terminal extensions showed naturally larger distances between these two helices already when interacting with canonical 9-mer peptides (P=0.002, Wilcoxon rank-sum test, Figure 4D). Using this distance to predict alleles accommodating C-terminal extensions among those with available crystal structures led to an AUC of 0.92. Of note, HLA-B51:01 is characterized by much higher frequency of 8-mer ligands compared to other HLA-I alleles (3, 5, 22). As for HLA-A01:01, we observed a trend in some samples (e.g., the mono-allelic cell line, see Table S2), but not in other samples (e.g., Melanoma/Mel_12, although most 10-mers are expected to come from HLA-A01:01 since HLA-B08:01 and HLA-C07:01 poorly bind 10-mers). This suggests that C-terminal extensions for HLA-A01:01, if present, show lower stability and may not be reproducible across samples. We finally point out that the clear patterns in both sequence and structural properties of HLA-I alleles displaying C-terminal extensions provides an additional and independent validation of our predictions in Figure 2 based only on HLA peptidomics data.

### Explicitly incorporating C-terminal extensions in HLA-I ligand predictors

C-terminal extensions have not been routinely investigated in previous studies and the training set of most existing HLA-I ligand predictors typically does not include them, even if recent version of NetMHC tools can mathematically handle them (23, 24). We therefore collected all C-terminally extended ligands uncovered in this work and retrained our predictor MixMHCpred (20) using multiple PWMs to model C-terminal extensions (25, 26) (Materials and Methods). To validate our new algorithm, we took advantage of the fact that HLA-A03:01 and HLA-A68:01 alleles were present in multiple datasets (see Table S1) and attempted to re-predict all the 10-mer peptides of these datasets, excluding data from the dataset used for testing in the training of our predictor (see Materials and Methods). As negative data, we included 4-fold decoy (i.e., 10-mers randomly selected from the human proteome) and computed both the Positive Predictive Value corresponding to the top 20% predictions (PPV, which in this case is also equivalent to the recall since the number of true positive is equal to the number of predictions) and the AUC (see Materials and Methods). Overall, we observed that explicitly modeling C-terminal extensions increased the performance compared to the previous version of our predictor (MixMHCpred1.0) (20), as well as other widely used HLA-I ligand predictors that did not include unbiased MS data in their training set (23, 24) (Figure 5A and Figure S6). The improvement came mainly from higher scores for ligands displaying the C-terminal extensions, as shown in Figure 5B for the HLA-A03:01 10-mer ligands isolated from a mono-allelic cell line (3) (P=1.0x10^-7^ for [R/K][A/F/I/L/M/Y]-COOH 10-mer peptides, Wilcoxon signed-rank test).

**Figure 5.**
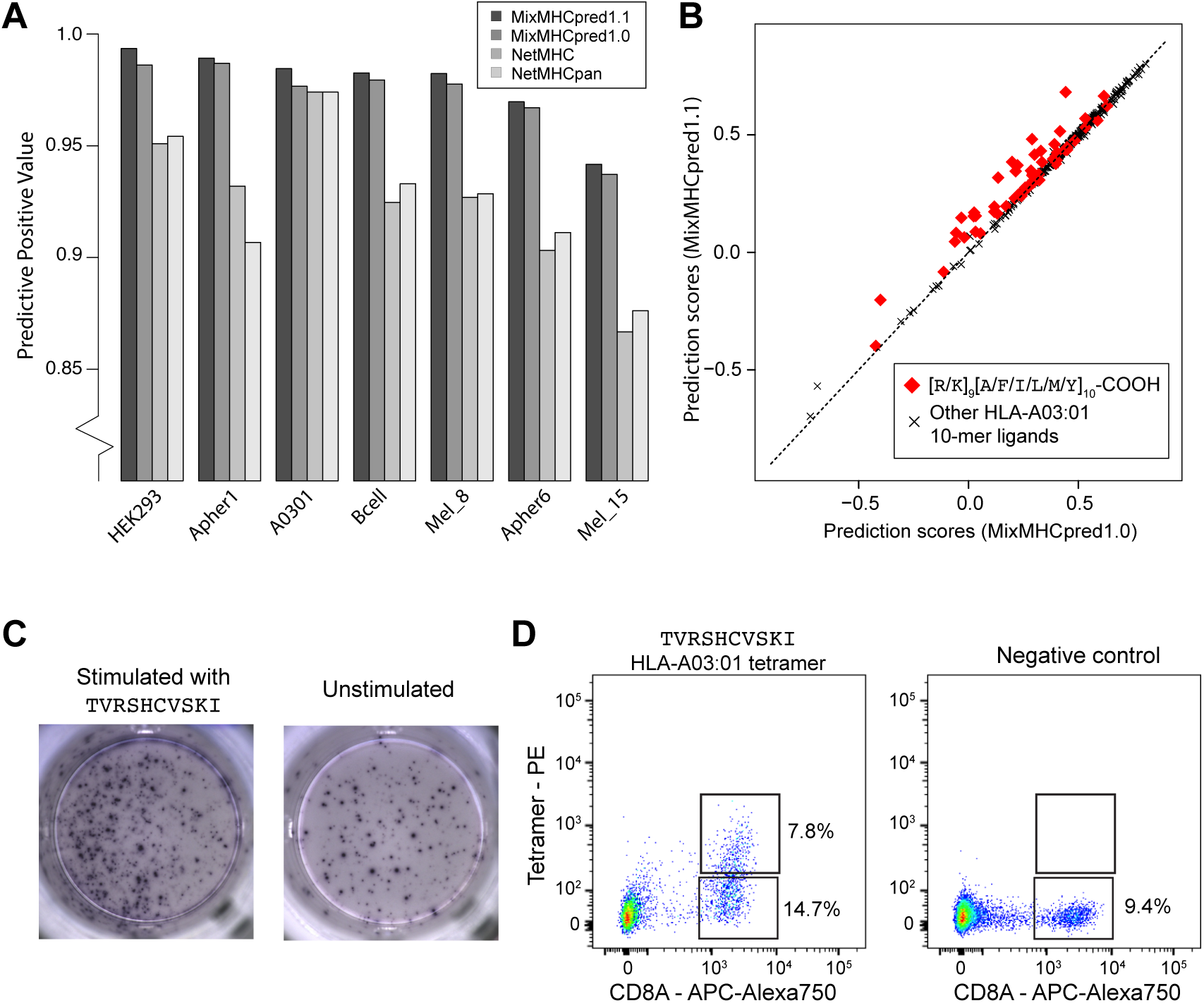
**A:** Benchmarking of the new version of our HLA-I ligand predictor (MixMHCpred1.1). The y-axis shows the Positive Predictive Value among the top 20% of the predictions. **B:** Analysis of binding prediction scores when explicitly modeling C-terminal extensions (MixMHCpred1.1) or not (MixMHCpred1.0) for the 10-mer HLA-A03:01 ligands from a mono-allelic cell line (3), as a function of the C-terminal amino acids (P9 and P10). **C:** IFNγ-ELISpot results obtained by stimulation with a CMV C-terminally extended 10-mer HLA-A03:01 peptide (`TVRSHCVSKI`; left) vs. no peptide (right) of a PBMC sample from a HLA-A03:01 and CMV seropositive healthy donor, previously stimulated for 12 days with the `TVRSHCVSKI` peptide. **D:** Multimer analysis of CD8 T cells from healthy donors recognizing the HLA-A03:01 restricted C-terminally extended 10-mer peptide `TVRSHCVSKI` (left) and the negative control (`RVRAYFYSKV`/HLA-A03:01 tetramer) for which we did not observe T cell recognition (right).

Almost no difference between MixMHCpred1.0 and MixMHCpred1.1 could be observed when testing our algorithm on other non-MS datasets used in previous benchmarking studies. However, we anticipate that C-terminal extensions are very rare in these datasets since most of the known HLA-I ligands had been first predicted with former versions of HLA-I ligand predictors that did not include C-terminal extensions. For instance, when analyzing IEDB data (27) for HLA-A03:01 ligands coming from earlier studies than those considered in this work, we could not see any statistical evidence of C-terminal extensions, suggesting that such ligands had not been tested in binding assays, or had been filtered in MS data.

### Identification of immunogenic C-terminally extended epitopes

To investigate the immunological relevance of HLA-I ligands displaying C-terminal extensions, we used our new predictor and scanned both human cancer testis antigens and viral proteomes with the motifs characterizing 10-mer C-terminal extensions for HLA-A03:01 and HLA-A68:01 (Materials and Methods). This analysis revealed many putative epitopes, including 10-mer peptides from the PRAME cancer testis antigen (`RLWGSIQSRY` for HLA-A03:01 and `ETLSITNCRL` for HLA-A68:01) as well as several CMV, EBV, HIV, HPV, Influenza and Yellow Fever peptides. To confirm that these peptides can indeed bind to HLA-A03:01 or HLA-A68:01, we measured the binding stability and found values falling within the range of CD8 T cell epitopes (see Table 1).

**Table 1:**
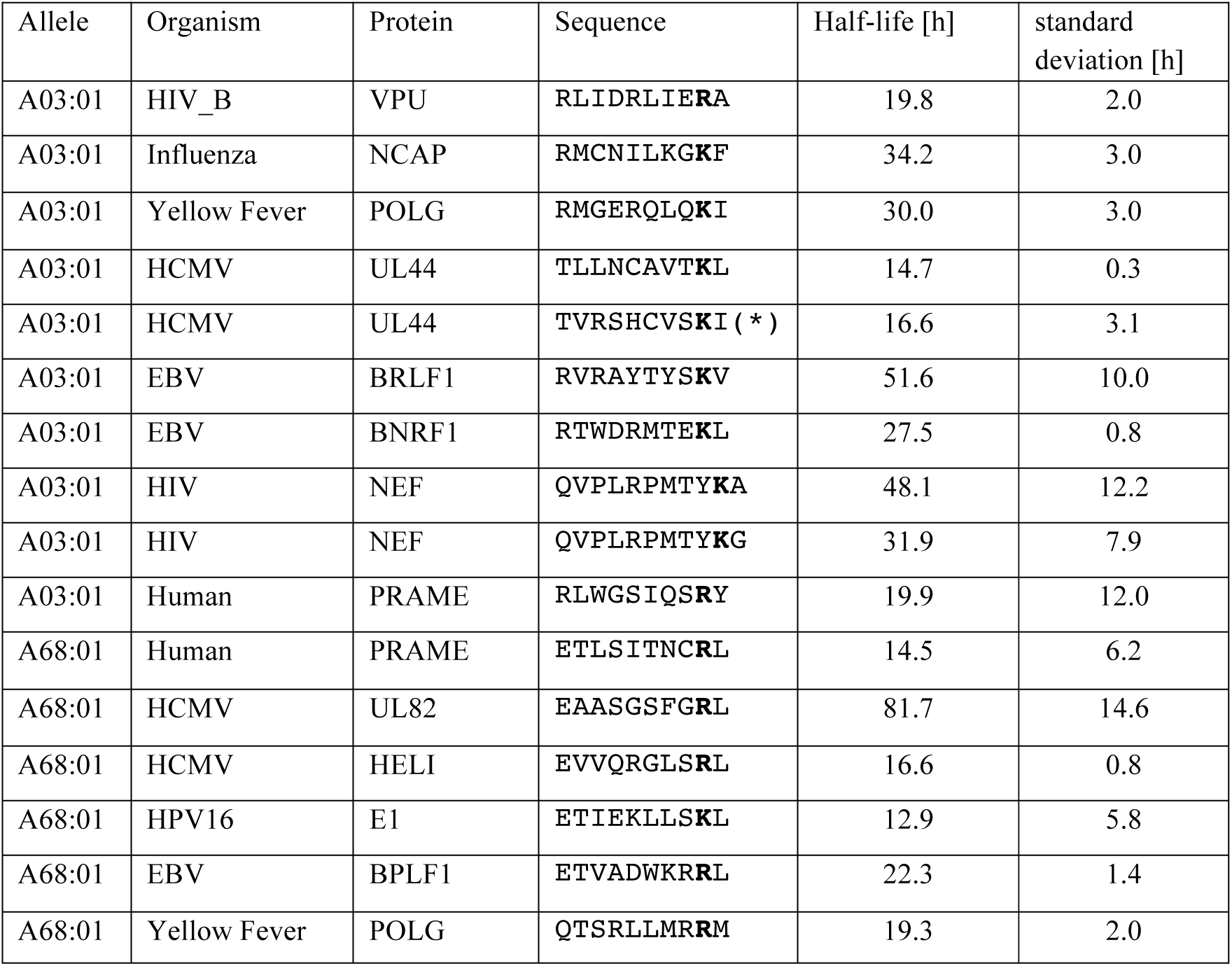
Binding stability (half-lives) of C-terminally extended HLA-I ligands from cancer testis antigens and viral proteins. The star indicates the peptide used in the ELISpot-IFNγ and multimer analyses of Figure 5C and 5D.

We then focused on one CMV peptide binding to HLA-A03:01 (star in Table 1). We stimulated a peripheral blood mononuclear cells (PBMCs) sample from a healthy donor (both HLA-A03:01 and CMV seropositive) with the 10-mer peptide for 12 days. Subsequently we observed cytokine production by IFNγ-ELISpot after re-challenge with the CMV 10-mer peptide for 16h (Figure 5C), suggesting that this epitope is potentially immunogenic in human. To determine whether CD8 T cells could directly recognize the C-terminally extended 10-mer peptide, we constructed HLA-A03:01 tetramers loaded with the 10-mer peptide. Analyzing CD8 T cells with such tetramers revealed a population of CD8 T cells that directly bound to the C-terminally extended 10-mer peptide/HLA-A03:01 complex (Figure 5D). This clearly indicates that the presence of C-terminal extensions in HLA-I ligands does not prevent TCR recognition and shows, for the first time to our knowledge, that C-terminally extended peptides can form *bona fide* CD8 T cell epitopes.

## Discussion

MS analysis of HLA-I ligands provides an unbiased view of HLA-I peptidomes that is not restricted by *a priori* assumptions on HLA-I binding specificity. MS-based approaches have therefore a high potential to identify new properties of HLA-I-peptide interactions. Here we capitalized on this premise to explore non-canonical binding modes of HLA-I ligands. Out of the 54 alleles considered in this work, we found clear statistical evidences of C-terminal extensions for eight of them and validated these predictions at the biochemical and structural level for the most frequent of these alleles in Caucasian populations.

In addition to providing statistical evidences of C-terminal extensions, our work enabled us to characterize new motifs describing this non-canonical binding mode. We observed that the C-terminal extensions are often characterized by the presence of hydrophobic amino acids, which could be structurally rationalized for HLA-A68:01. We also note that, for three out of eight alleles (i.e., HLA-A03:01, HLA-A31:01 and HLA-A68:01), positively charged amino acids are found at the anchor residue filling the F pocket. This positively charged residue interacts with D77 and D116 in our HLA-A68:01 structure, which is consistent with the observation that Aspartate is preferentially observed at these positions in alleles accommodating C-terminal extensions (Figure 4C). The clearly distinct binding specificity between the anchor residue and the C-terminal extension makes these cases especially amenable for the statistical model that we developed to analyze HLA peptidomics data. However, charged residues at the C-terminal anchor positions only characterize a minority of HLA-I alleles, and the majority of alleles show preference for hydrophobic residues. In these cases, and assuming that the preference for hydrophobic amino acids at the C-terminal extension is conserved, sequenced-based algorithms cannot unambiguously determine whether a 10-mer peptide with 2 hydrophobic amino acids at the last two positions follows the bulge or the C-terminal extension binding mode. Moreover, we anticipate that competition with high affinity (mainly 9-mer) ligands can mask cases of less frequent C-terminal extensions in HLA peptidomics data. Therefore our estimates of the number of alleles that accommodate C-terminal extensions correspond to lower bounds and we cannot exclude that this non-canonical binding mode may be observed in other alleles. In particular, we cannot exclude that some C-terminal extensions among HLA-A02:01 ligands may be present in our data with two hydrophobic residues at the last positions. Nevertheless, the smaller distance between the two alpha helices of HLA-A02:01 (Figure 4D) suggests that C-terminal extensions for this allele should be rarer, or involve other processing and presentation mechanisms, like cross-presentation (10, 11). Along this line, we note that statistically significant C-terminal extensions are identified by our algorithm in the set of *Toxoplasma gondii* HLA-A02:01 ligands (10) (see Table S5), confirming that our approach is able to capture previous evidences of C-terminal extensions.

Finally, we stress that contaminations from co-eluting or wrongly identified peptides, as well as challenges in aligning small peptides or in the deconvolution of pooled HLA peptidomics datasets, can easily result in cases that look like N- or C-terminal extensions by chance. This is the reason why we developed the statistical framework described in this work and extensively tested the robustness of our predictions with respect to noise in the data.

Despite these inherent limitations of sequence-based approaches, it is likely that several alleles show very few if any C-terminal extensions. For instance, six alleles from our list showed specificity at the last anchor residue that is not restricted to hydrophobic amino acids (mainly tyrosine, see Figure S7) but we did not see any trend of C-terminal extensions among their 10-mer ligands, both in pooled and in mono-allelic HLA peptidomics data (see Table S2). Among them, all that had available X-ray data displayed a smaller distance between the two alpha helices (D_80-134_), which is consistent with our observation in Figure 4D.

Our model also incorporated the possibility to have N-terminal extensions but we did not find any such event, although N-terminal extensions have been recently reported (12, 13). Inspection of existing structures of HLA-I molecules in complex with 9-mer peptides shows that the C-terminal carboxyl group of 9-mer HLA-I ligands is often partly solvent exposed. Conversely, the N-terminal amide group points toward the binding site and is significantly less surface accessible in most HLA-I structures. This likely explains why N-terminal extensions appear to be much less frequent, although we cannot exclude that our approach may miss some N-terminal extensions if the specificity at P2 is the same as at P3 in the N-terminally extended ligands. Interestingly, this appears to be the case for N-terminal extensions reported for HLA-B57:01 and may explain why these extensions could not be easily detected with sequence-based approaches only and have been first identified by X-ray crystallography (12).

Our demonstration of direct recognition of C-terminally extended HLA-I ligands by human CD8 T cells (Figure 5D) shows that C-terminal extensions are compatible with TCR binding. C-terminal extensions may further play a role in the binding of KIR proteins, which are known to recognize mainly amino acids surrounding the C-terminus of canonical HLA-I ligands (12, 28, 29). As such, we anticipate that inclusion of C-terminal extensions the new version of our HLA-I ligands predictor for alleles displaying frequent C-terminal extensions may help uncovering new CD8 T cell epitopes in viruses or tumors, as well as potential targets of NK cells.

Overall, our results reveal frequent C-terminal extensions in at least eight out of the 54 HLA-I alleles analyzed in this study and highlight the power of unbiased HLA peptidomics data together with new algorithms to unravel novel properties of HLA-I molecules. Our demonstration of direct T cell recognition of such epitopes suggests that C-terminal extensions may be clinically relevant for infectious diseases or cancer immunotherapy.

## Materials and Methods

### Collection of HLA peptidomics data

HLA peptidomics data used in this study came from seven different studies (3, 15–20) (see Table S1). In all these studies, peptides were eluted from class I specific antibody purified HLA molecules. None of these data had been filtered with existing predictors, and therefore all had the potential to reveal non-canonical ligands. We only included MS samples generated with < 1% false discovery rate, with available HLA-I typing information and in which HLA-I motifs could be clearly annotated by our motif deconvolution approach for pooled HLA peptidomics data (4, 20). In total, our dataset comprises 43 samples covering 54 different HLA-I alleles (see Table S1) for a total of 109,953 unique peptides (9- to 12-mers).

### Analysis of non-canonical binding modes

#### Analysis of 9-mer HLA peptidomes

To analyze non-canonical binding modes in pooled HLA peptidomics studies that include ligands from up to six different HLA-I molecules, we developed the following pipeline (Figure 1B and 1C). We first characterized the different motifs of the 9-mer HLA peptidomes using our recent motif deconvolution algorithm (4). This algorithm has the major advantage of not depending on *a priori* knowledge of HLA-I binding specificity, and therefore can be applied even in the presence of poorly characterized HLA-I alleles. Each 9-mer peptide was then assigned to its corresponding motif and PWMs were built for each group of peptides, including random counts based on BLOSUM62 (20).

#### Prediction of N- and C-terminal extensions

To distinguish between the three different models describing longer peptides (i.e. bulge, N-terminal extensions, C-terminal extensions), we developed the algorithm illustrated in Figure 1C. We first reasoned that longer peptides following the bulge model should have conserved binding specificity around the anchor residues at the second and last positions. We therefore modeled these peptides with the binding specificity of 9-mers at the first three and last two amino acids, and unspecific position in the middle. In the absence of a priori information about the amino acid preferences at terminal extensions, we modeled them as with one unspecific position at the N- or C-terminus. To enable meaningful comparison between the scores of the bulge and the terminal extension models, we did not impose any constrain at middle positions in the N- or C-terminal extension models. Each longer peptide (i.e., *L*-mer, with *L* equal to 10, 11 or 12) was then scored with all models (i.e. bulge, N- or C-terminal extensions) derived from each motif identified in the 9-mer HLA peptidome. Scores for the different models were computed as: 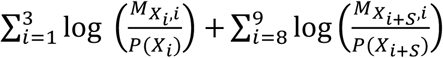 for the bulge model; 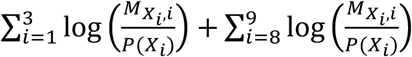 for the C-terminal extension model and 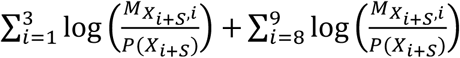 for the N-terminal extension model, where M stands for the 20x9 PWM derived from 9-mer ligands for a given allele, X stands for the sequence of a *L*-mer peptide, *S* equals to *L*-9, and *P(X*_*i*_) stands for the frequency of amino acid *X*_*i*_ in the human proteome. Peptides were then assigned to one model and one allele if their score with this allele and this model was higher than a threshold T_1_ and no other score for all the other possible models of any allele was larger than the largest score minus T_2_. Here we chose values of T_1_ and T_2_ equal to 2.0, since over 95% of the longer peptides found in our HLA peptidomics datasets had a score larger than this value for at least one allele and one model.

Alleles with anchor residues at positions 4 to 7 (i.e., HLA-B08:01, HLA-B14:01 and HLA-B14:02) were excluded, as they display non-conserved binding motifs between 9- and 10-mers and much less 10-mer ligands (4).

#### Null model

To account for possible noise in MS data and focus on predictions that showed statistical significance, we developed a null model representing the expected HLA peptidome from the bulge model, and described here for the case of 10-mer ligands. Starting with a list 100’000 10-mer peptides randomly selected from the human proteome, we selected those that passed the threshold T_1_ for the bulge model of at least one motif. The list of peptides was further randomly filtered so as to have the same number of peptides assigned to each allele as in the actual 10-mers HLA peptidomics data. We then re-predicted these peptides using all three models for each motif (Figure 1C). This enabled us to assess how many 10-mer peptides generated from the bulge model of each allele could by chance be assigned to the N- or C-terminal extensions of any allele when considering the three models. The simulations were repeated 100 times to derive a Z-score. Only alleles with at least 50 10-mer peptides assigned to them, at least 10 peptides assigned to the C- or N-terminal extension model and with a Z-score larger than 2 were included in our predictions. Similar predictions were obtained using values ranging between 1.5 and 2.5 for the threshold T_1_, or using a different threshold T_1_ for each motif given by the score corresponding to the top 2% predictions in a large set of 100’000 10-mer peptides randomly selected from the human proteome (Table S6).

The fractions of C-terminal extensions shown in Figure 2B and 2D were computed as the number of peptides unambiguously assigned to the C-terminal extension mode of a given allele, divided by the number of peptides assigned to any of the three models for the same allele.

Sequence logos were generated with the LOLA software (http://baderlab.org/Software/LOLA).

### In vitro binding assays

All peptides shown in Figure 3, Figure S2 and Table 1 were synthesized with free N and C-termini (1mg of each peptide, > 80% purity). Peptides were incubated separately with denaturated HLA alleles refolded by dilution in the presence of biotinylated beta-2 microglobulin proteins at temperature T=4°C for 48 hours. The solution was then incubated at 37°C. Samples were retrieved at time t=0h, 8h, 24h, 48h and t=72h. Stable complexes indicating interactions between HLA-I molecules and the peptides were detected by ELISA. Signals for 9-mer ligands at time t=0h were used for renormalization in Figure 3 and Figure S2, while negative controls consisted of absence of peptides. Half-lives (Table 1) were computed as ln(2)/k_off,_ where k_off_ were determined by fitting exponential curves to the light intensity values obtained by ELISA, after removing the background signal (i.e., no peptides). Two independent replicates were performed for each measurement.

### Expression and purification of HLA-A68:01 and β2M

The heavy chain HLA-A68:01 and the light chain β2M were purified from inclusion-bodies (30) with some small modifications. Briefly, recombinant expression plasmids were transformed into BL21 (DE3) (HLA-A68:01) or XA90 strain (β2M) bacteria. The cells were grown overnight in LB (Luria-Bertani medium) twice concentrated supplemented with 50 µg ml^-1^ kanamycin or 100 µg ml^-1^ ampicillin respectively at 37 °C. One litre of pre-warmed LB was inoculated with 10 ml of the overnight culture and was incubated at 37 °C. At an OD_600 nm_ of 0.6, expression was induced for 8 h at 37 °C with 1.0 mM isopropyl-β-D-thiogalactopyranoside (IPTG). Cells were then harvested by centrifugation (8700x*g*, 15 min, 4 °C, Beckman Coulter Avanti J-20 XP centrifuge), and then re-suspended in lysis buffer (10 mM TRIS-HCl, pH 8.0 at 20 °C complemented with 100 µg ml^-1^ Lysozyme (Sigma Aldrich), 250 units of Benzonase (Novagen), 1 mM EDTA and 1:1000 (v/v) Protease Inhibitor Cocktail III (Calbiochem)). After 20 min of continuous rocking at 22 °C, cells were lysed 3 times at 4 °C using a Basic Z-Model Cell Disrupter (Constant Systems Ltd, UK). Inclusion-bodies were isolated by centrifugation (16000x*g* for 1 h at 4 °C, JA 25.50 rotor, on a Beckman Coulter Avanti J-20 XP centrifuge). The supernatant was decanted and the contaminated material was removed by scrapping the outer rings (viscous and dark coloured) leaving the more compact and lighter coloured inner ring (containing the protein of interest) intact. The pellet was then fully dispersed in 20 ml of washing buffer (10 mM TRIS-HCl, pH 8.0 at 20 °C) complemented with 1:1000 (v/v) Protease Inhibitor Cocktail III (Calbiochem) and centrifuged again for 10 min (16000x*g* at 4 °C, JA 25.50 rotor, on a Beckman Coulter Avanti J-20 XP centrifuge). The washing/scrapping steps were repeated 5 times (until the outer rings vanished). The pellet containing the recombinant protein was then dissolved in 20 ml of solubilisation buffer (100 mM TRIS-HCl, pH 8.0 at 20 °C, 8 M urea). Insoluble material was precipitated by centrifugation (16000x*g* for 1 h at 4 °C, JA 25.50 rotor, on a Beckman Coulter Avanti J-20 XP centrifuge). Solubilized HLA-A68:01 heavy chain was immediately flash frozen in liquid nitrogen and stored at -80 °C. The recombinant β2M protein in urea was refolded by dialysis against 10 mM TRIS-HCl pH7.0 using SnakeSkin^®^ Dialysis tubing (3.500 MWCO, Thermo Scientific) and purified by ion exchange on Hi-Trap Q HP (5 ml, GE Healthcare) column, with a linear gradient from 0 to 100 mM NaCl. Fractions containing pure β2M were dialysed overnight against water, concentrated with a 10 MWCO concentrator (Amicon^®^ Ultra, MILLIPORE) to 2 mg ml^-1^, flash frozen in liquid nitrogen and stored at -80 °C. HLA:β2M:peptide complex assembly

The protein complex was reconstituted by dilution of the denatured HLA-A68:01 heavy chain (3 μM) and β2M (6 μM) in presence of the peptide ([H]-E-T-S-P-L-T-A-E-K-L-[OH], 10 μM) into 200 ml of refolding buffer (100 mM TRIS-HCl, pH 8.0 at 20 °C, 400 mM L-Arginine HCl (Sigma Aldrich), 2 mM EDTA, 5 mM reduced L-glutathione (Sigma Aldrich), 0.5 mM oxidized L-glutathione (Sigma Aldrich) and with 1:1000 (v/v) Protease Inhibitor Cocktail III (Calbiochem)). The refolding mixture was incubated at 10 °C during 36 h under constant stirring. Every 12 h another batch of the denatured HLA-A68:01 heavy chain (3 μM) was added to the mix. The 200 ml were then concentrated to 5 ml using a 3 MWCO concentrator (Amicon^®^ Ultra, MILLIPORE). The concentrated protein mixture was further submitted to exclusion size chromatography (HiLoad™ 16/60 Superdex™ 75 prep grade, on Äkta Pure GE Healthcare) and each peak collected was submitted to SDS-PAGE gel and mass spectrometry (Agilent 6530 QTOF (Agilent Technologies Inc. - Palo Alto, CA)) to assess the simultaneous presence of the 3 components of the complex. Fractions of interest were then stored at 4°C until further use.

### Crystallization

Prior to crystallization, the buffer of the protein complex was exchanged to 25 mM MES pH6.5 at 20 °C and 150 mM NaCl on a Superdex™ 200 Increase 10/300 GL column using an Äkta Pure system (GE Healthcare). The complex was then concentrated to 10.96 mg ml^-1^ final using a 3 kDa MWCO concentrator (Amicon^®^ Ultra, MILLIPORE). Aliquots of the complex were set up for crystallization using a mosquito^®^ crystallization robot (TTP Labtech, Royston UK). Coarse screens were typically setup onto Greiner 3-well plates using three different drop ratios of precipitant to protein per condition (100+50 nl, 75+75 nl and 50+100 nl). Crystallization was carried out using the sitting drop vapor diffusion method at 4 °C. Crystals were grown by mixing 100 nl of the protein complex (10.96 mg/ml) with 50 nl of reservoir solution containing 0.1 M HEPES pH 7.5, 12 % PEG3350, 0.005 M CoCl_2_, 0.005 M NiCl_2_, 0.005 M CdCl_2_ and 0.005 M MgCl_2_. Diffraction quality crystals grew within a few days.

### Data Collection and Structure Refinement

Crystals were cryo-protected using the well solution supplemented with additional ethylene glycol and were flash frozen in liquid nitrogen. Data were collected at Diamond beamline I24 on a Pilatus3 6M detector at a wavelength of 0.96864 Å. Indexing and integration was carried out using XDS (31) and scaling was performed with SCALA (32). Initial phases were calculated by molecular replacement with PHASER (33) using a model of HLA/β2M peptide complex (PDB:5T6X). Initial models were built by ARP/wARP (34) followed by manual building in COOT (35). Refinement was carried out in REFMAC5 (36). Thermal motions were analyzed using TLSMD (37) and hydrogen atoms were included in late refinement cycles. Data collection and refinement statistics can be found in Table S7. The model and structure factors have been deposited with PDB accession code: 6EI2 (see Preliminary_Validation_report.pdf).

### HLA-I sequence and structure analysis

For all 54 alleles considered in this work, the sequence of the peptide binding domains were retrieved from IMGT database (38) and aligned with MUSCLE (39). To investigate the molecular determinants of C-terminal extensions, we selected all amino acids surrounding the F pocket (75-81, 84, 114-118, 142-147, following residue numbering in HLA-I structures). The eight positions displaying the largest changes in amino acid frequencies (Jensen-Shannon divergence larger than 0.1) are shown in Figure 4C. Using 20-dimensional encoding for each amino acid, we trained a logistic regression model based on *glmnet* package in R (40). Four-fold cross-validation was performed and repeated 10 times for different random seeds to compute AUCs.

Available HLA-I crystal structures were then collected for alleles considered in this work. In total, 20 alleles among those with available HLA peptidomics data had experimental X-ray crystal structures. For each allele, a reference structure was selected by prioritizing complexes with 9-mer ligands and highest resolution (see Table S4). In each structure, the distance between the C-alpha of residues 80 and 143 was computed (Figure 4D and Table S4). The AUC of a predictor using as input feature this distance was 0.92, which is comparable to the AUC of the predictor based on HLA-I sequences when considering only the 20 HLA-I alleles with available crystal structures (AUC=0.93, three-fold cross-validation).

### HLA-I ligand predictors explicitly incorporating C-terminal extensions

For each allele displaying C-terminal extensions, we retrained our predictor based on HLA peptidomics data modeling C-terminal extensions as multiple motifs. In practice, all 10-mer ligands predicted to follow the C-terminal extension mode of each allele were treated separately and a PWM (*M*^*C*^) was built with them (20). All other 10-mer ligands (non-C-terminal extensions) were used to build another PWM describing the bulge model (*M*^*b*^). For alleles with C-terminal extensions, the score of a 10-mer peptide X=(X_1_, … X_10_) with this predictor is then computed as:

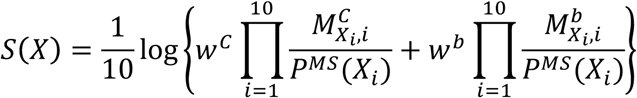

where *w*^*C*^, respectively *w*^*b*^, stands for the fraction of ligands predicted to follow C-terminal extensions, respectively bulges, and *P*^*MS*^ stands for the background distribution of amino acids in HLA peptidomics data (see (20)). For alleles without C-terminal extensions, the predictions did not change compared to the previous version of our predictor (20).

Benchmarking of the algorithm was done using HLA-A03:01 and/or HLA-A68:01 positive samples, since these are the two HLA-I alleles displaying C-terminal extensions in multiple samples in our collection of HLA peptidomics studies. Careful cross-validation was performed, where for each sample used as testing set, the predictor was trained only on the data from the other samples. To this end, we further excluded from our analysis samples where some other HLA-I alleles did not appear in other samples (i.e., Fibroblast and RA957). For each sample, 4-fold excess of random peptides from the human proteome were added as negatives and all peptides were ranked with our new predictor, using for each peptide the highest score across the different HLA-I alleles present in the sample. The fraction of positives found in the top 20% of the predictions as well as the AUC were then computed (Figure 5A and Figure S6). The performance of the predictor explicitly modeling C-terminal extensions (MixMHCpred1.1) was compared with the former version of MixMHCpred (v1.0, based on a single PWM for each allele) (20), NetMHC4.0 (24) and NetMHCpan3.0 (23).

### Human PBMC, ELISpot assays and Multimer analysis

Healthy volunteers donated circulating leukocytes according to the standards of the Blood Transfusion Center in Epalinges, Switzerland (Service Vaudois de Transfusion Sanguine). Samples from patients were obtained under written informed consent following study protocol approval by the Human Research Ethics Committee of the Canton de Vaud (Switzerland).

PBMCs were peptide stimulated in vitro for 12 days in vitro in presence of 100 U/ml IL-2. Subsequently, the Elispot was performed using the ELISpot^PRO^ kit for Human IFN-γ from MABTECH (3420-2APT-10), following the standard supplier instructions. 100’000 cells per well were re-challenged with the peptide for 16h. The spots were analysed by the iSpot Robot ELISpot reader (AutoImmun Diagnostika GMBH).

Monomeric HLA-A03:01/`TVRSHCVSKI` complexes were prepared by refolding procedures using heavy chain HLA-A03:01 and the light chain β2M, as described before. The heavy chain contained added BSP (BirA enzyme Substrate Peptide). The enzymatic biotinylation of HLA-A3:01-BSP was performed over night at 25°C with ATP, biotin and the biotin ligase Bir A. Multimer complexes were prepared by mixing biotinylated HLA-A03:01/`TVRSHCVSKI` monomers with PE-labeled streptavidin (Invitrogen).

CD8 T cells of the CMV 10-mer peptide stimulated PBMC after 12 days in vitro stimulation were stained with a PE labeled HLA-A03:01/`TVRSHCVSKI` multimer and co-stained with anti-CD8 antibody (BC A94683) and DAPI for dead cell exclusion. Multimer+ CD8+ T cells were analysed at the BD ARIA III instrument equipped with the FACS Diva software. The analysis was performed with the FlowJo software (FLOWJO.LLC). The negative control consists of an EBV-derived peptides (`RVRAYFYSKV`)/HLA-A03:01 tetramer and a PBMC sample from another HLA-A03:01 and EBV seropositive healthy donor, for which we did not observe any recognition by CD8 T cells.

## Code availability

The code for C- and N-terminal extensions (MHCpExt) and the new version of the predictor (MixMHCpred1.1) are available at https://github.com/GfellerLab.

## Acknowledgment

We thank Julien Racle and Santiago Carmona for a critical reading of the manuscript. DG acknowledges the financial support of CADMOS. We thank Anne Wilson and her collaborators of the flow cytometry facility, and the members of our laboratories for their support. Peptides were synthesized at the Protein and Peptide Chemistry Facility. The authors are grateful for support received by the SGC, a registered charity (number 1097737) that receives funds from AbbVie, Bayer Pharma AG, Boehringer Ingelheim, Canada Foundation for Innovation, Eshelman Institute for Innovation, Genome Canada, Innovative Medicines Initiative (EU/EFPIA) (ULTRA-DD Grant 115766), Janssen, Merck & Co., Novartis Pharma AG, Ontario Ministry of Economic Development and Innovation, Pfizer, S~ao Paulo Research Foundation-FAPESP, Takeda, and the Wellcome Trust (092809/Z/10/Z). P.F., and S.P. are supported by a Wellcome Career Development Fellowship (095751/Z/11/Z). We also thank Diamond Light Source for access to beamline I24 (proposal number mx15433), which contributed to the results presented here. Computations were performed at the Vital-IT (http://www.vital-it.ch) Center for high-performance computing of the Swiss Institute of Bioinformatics.

## Contributions

D.G. designed the study, developed the algorithms, analyzed the data and wrote the manuscript. P.G., P.B., N.M. and J.S. performed the experiments. S.P and P.F performed the X-ray crystallography. D.E.S, and G.C. contributed with reagents, M.B.-S., and D.E.S provided feedback and manuscript revision.

## Supplementary Material

**Figure S1:**
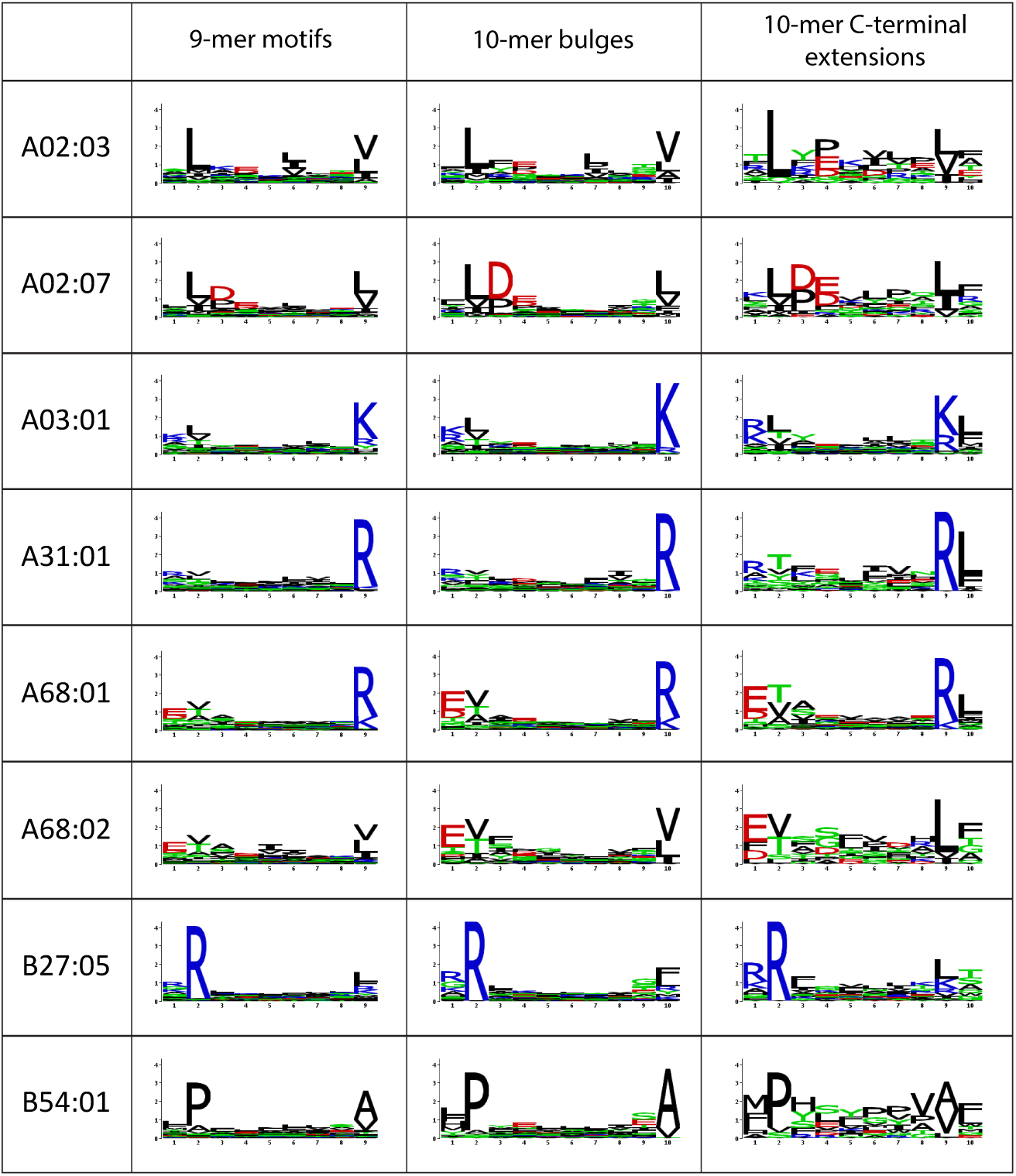
Comparison between 9-mer motifs, 10-mer motifs from peptides predicted to follow the bulge model and 10-mer motifs from peptides predicted to display C-terminal extensions for the eight alleles with such extensions. For HLA-A03:01 and HLA-A68:01, peptides have been pooled from the different studies where C-terminal extensions were identified to generate the logos.

**Figure S2:**
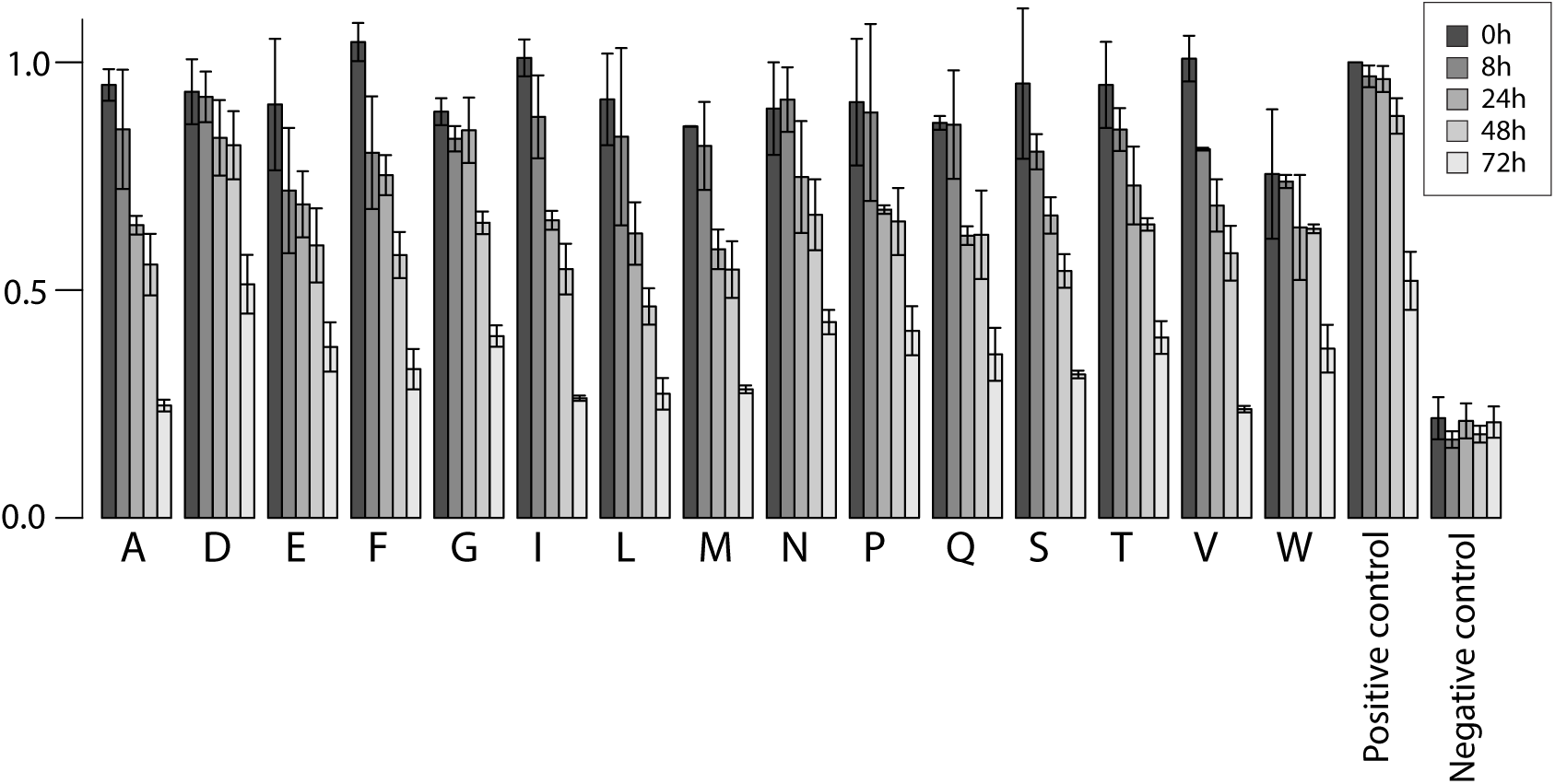
Stability analysis of 11-mers built from the C-terminally extended 10-mer KLAYTLLNKL HLA-A03:01 ligand (see Figure 3) with all possible C-terminal amino acids not compatible with the specificity of the second anchor for HLA-A03:01 (i.e., KLAYTLLNKL[A/D/E/F/G/I/L/M/N/P/Q/S/T/V/W]). ILRGSVAHK was used as positive control. Negative control consists of absence of peptide.

**Figure S3:**
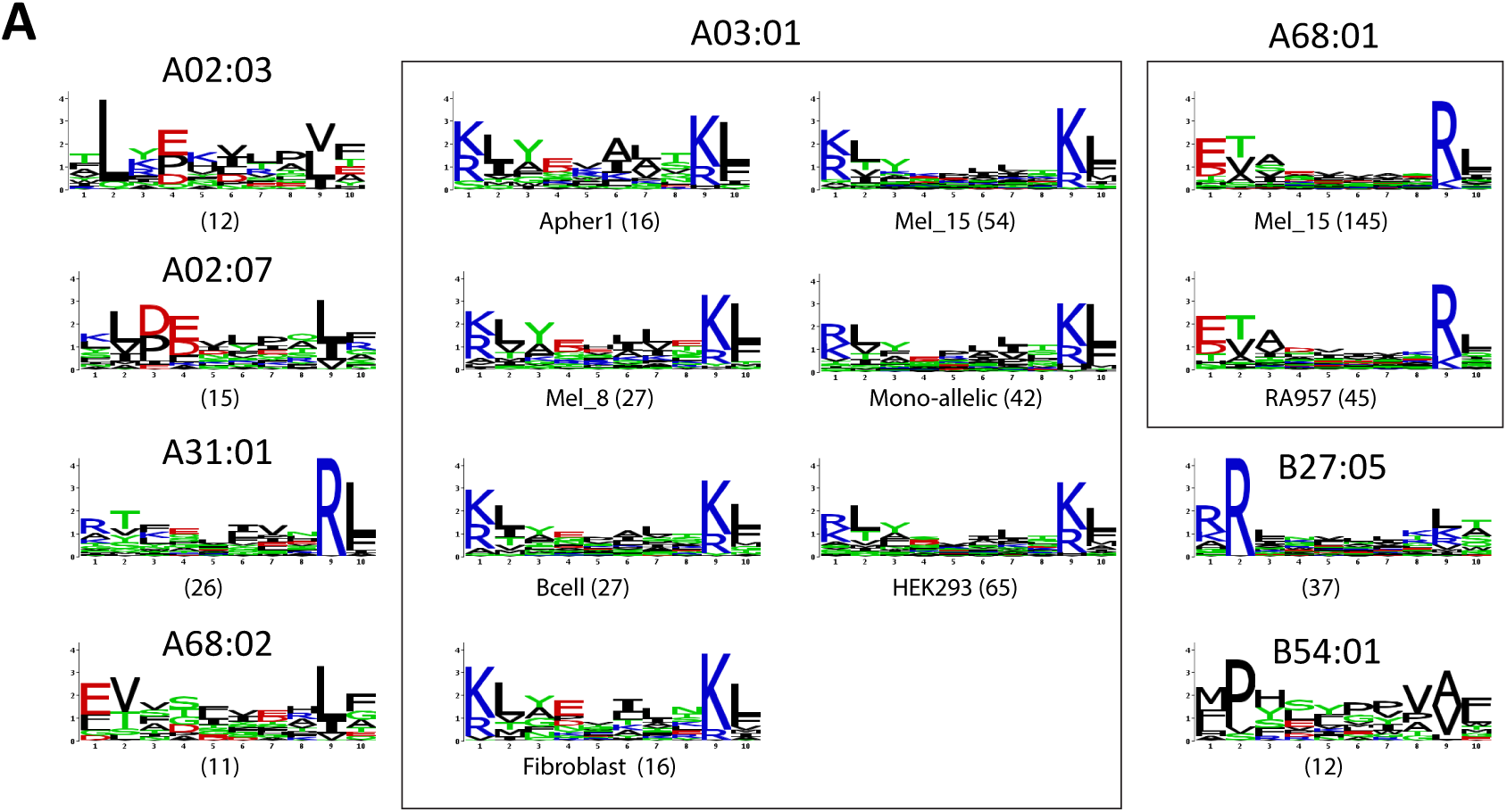
Predictions of C-terminal extensions in the presence of 5% of noise in all HLA peptidomics datasets.

**Figure S4:**
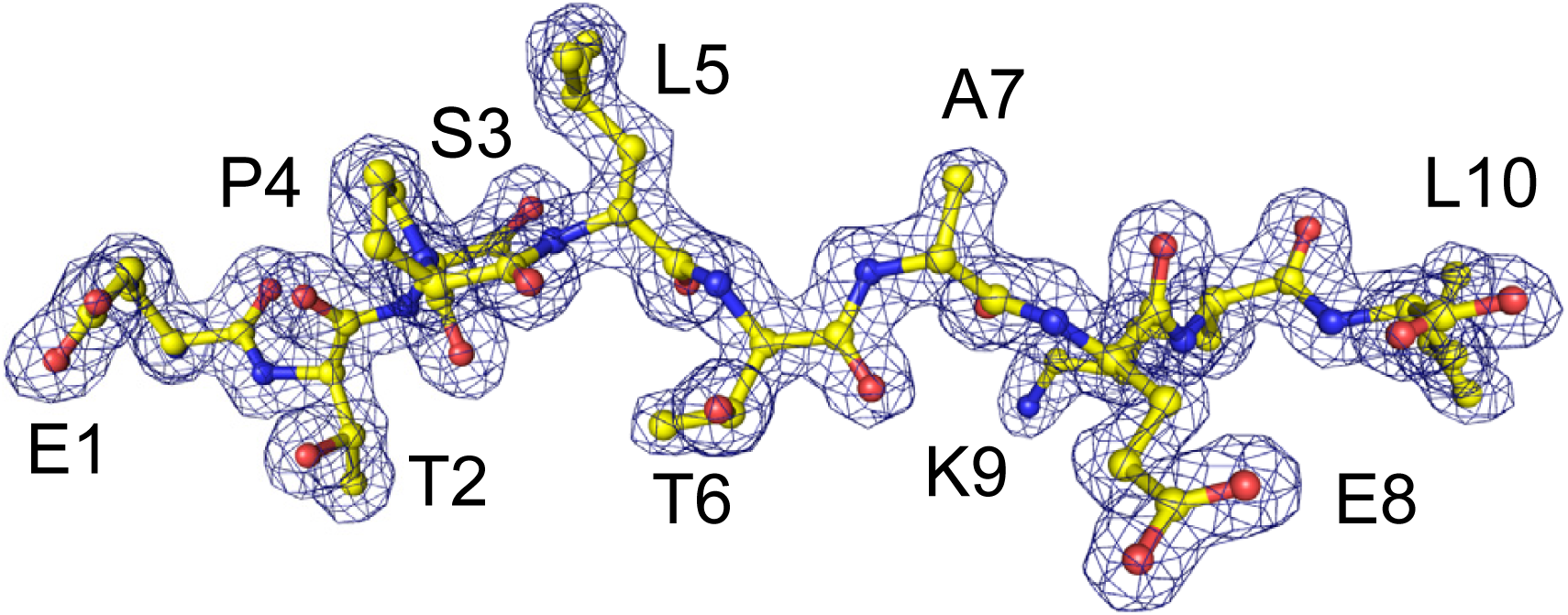
Map of the peptide's experimental electron density in the new X-ray structure shown in Figure 4A (Resolution 1.6Å).

**Figure S5:**
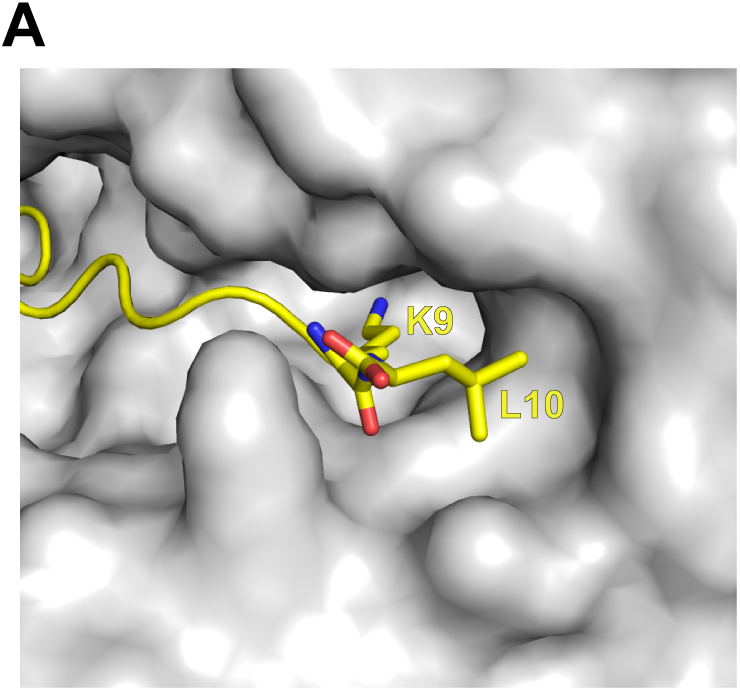
Surface view of the new structure of HLA-A68:01 in complex with the C-terminally extended peptide (ETSPLTAEKL).

**Figure S6:**
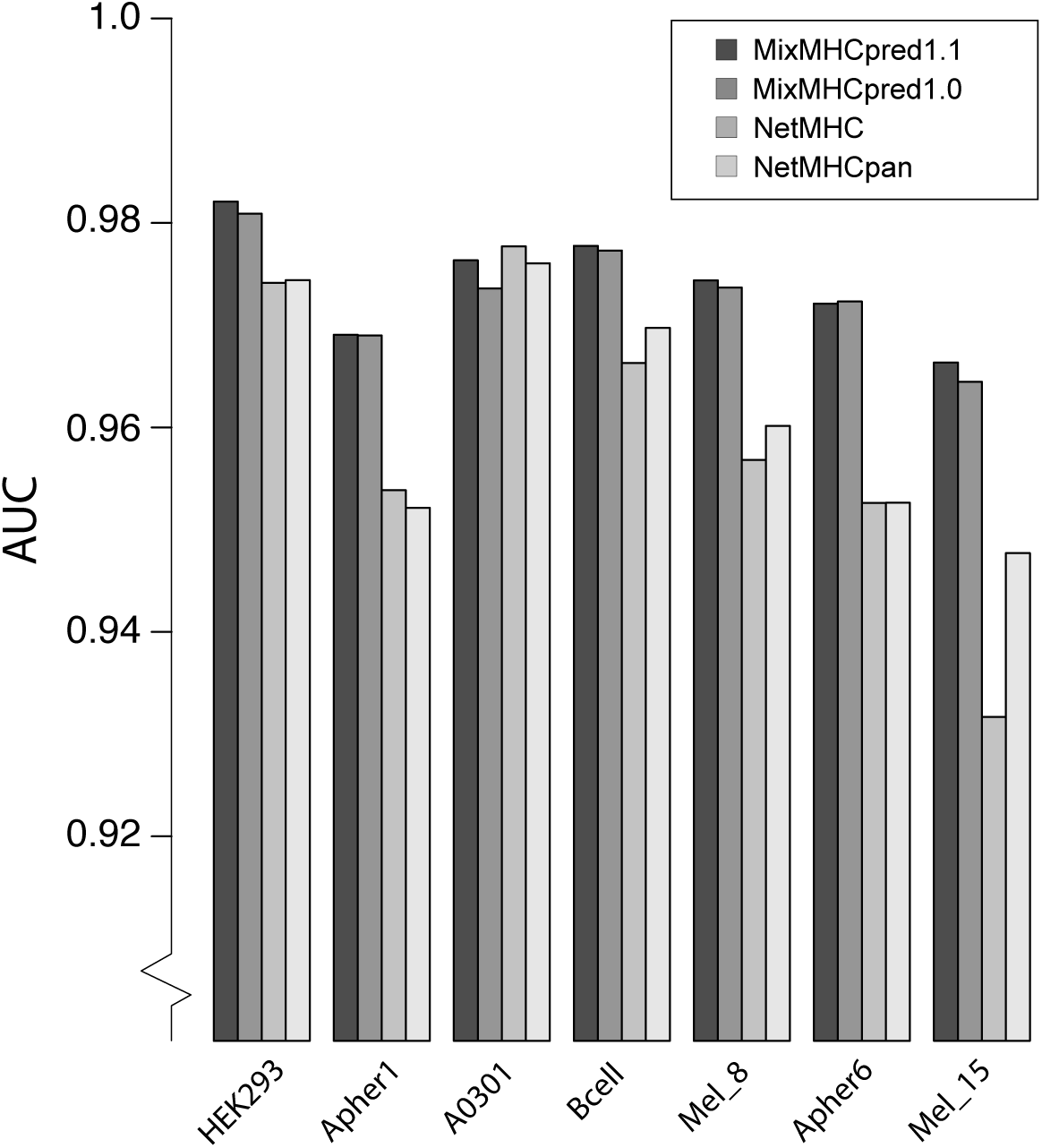
AUC values for different predictors when re-predicting HLA peptidomics data from samples containing HLA-A03:01 or HLA-A68:01 alleles (same cross-validation study as in Figure 5A).

**Figure S7:**
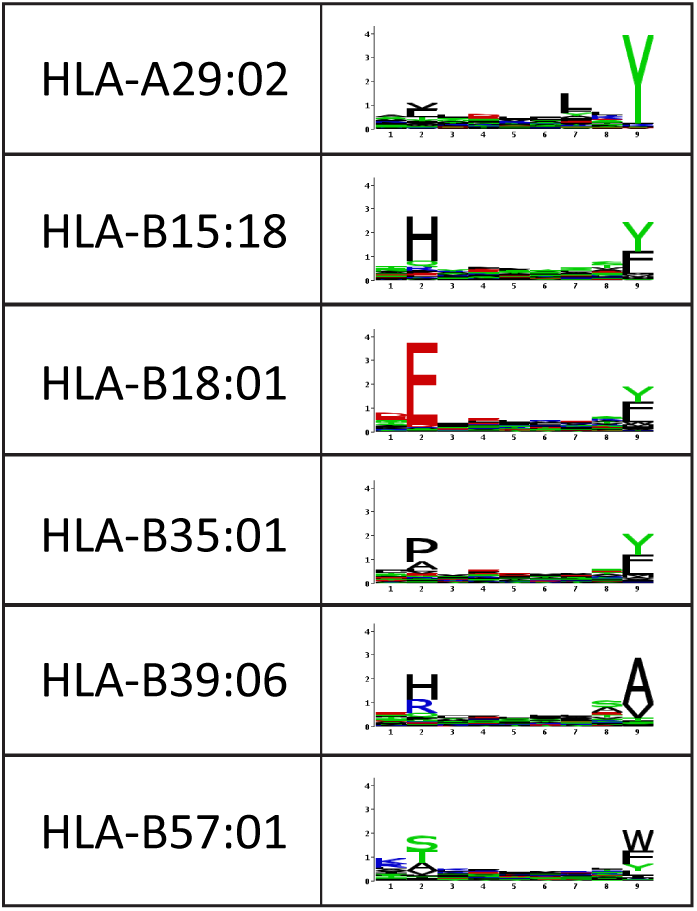
Examples of alleles, together with the 9-mer motifs, for which no C-terminal extensions were observed and that do not only show preference for hydrophobic amino acids at the C-terminal anchor residue.

**Table S1:** List of HLA peptidomics samples considered in this work with HLA typing information. Melanoma (15); MCP (16); Mommen (18); Ritz (17); Lausanne_2016 (20); Abelin (3); Hilton (19).

**Table S2:** Results of predictions of C-terminal extensions for 10-mer and 11-mer peptides. The first two columns indicate the sample of origin. Column 3 indicates the allele to which one 9-mer motif was annotated (20). Column 4 shows the total number of 10-mers with one-residue long C-terminal extension, respectively 11-mers with two-residue long C-terminal extensions. Column 5 shows the number of C-terminal extensions predicted for the allele in Column 3. Column 6 and 7 show the expected number and standard deviation over 100 realizations of the 10-mer peptidome assuming only bulges (see Materials and Methods). Column 8 shows the Z-score. NA stands for either motifs that could not be annotated, or cases where the number of 10-/11-mer ligands associated with the allele in column 3 did not pass our threshold (mainly B08:01 and HLA-C alleles which poorly bind to longer peptides).

**Table S3:** List of 10-/11-mer peptides predicted to display C-terminal extensions for samples and alleles for which the number of C-terminal extensions passed our thresholds. These peptides were used in Figure 2 and pooled together to define the final motifs for each allele (Figure S1, last column).

**Table S4:**
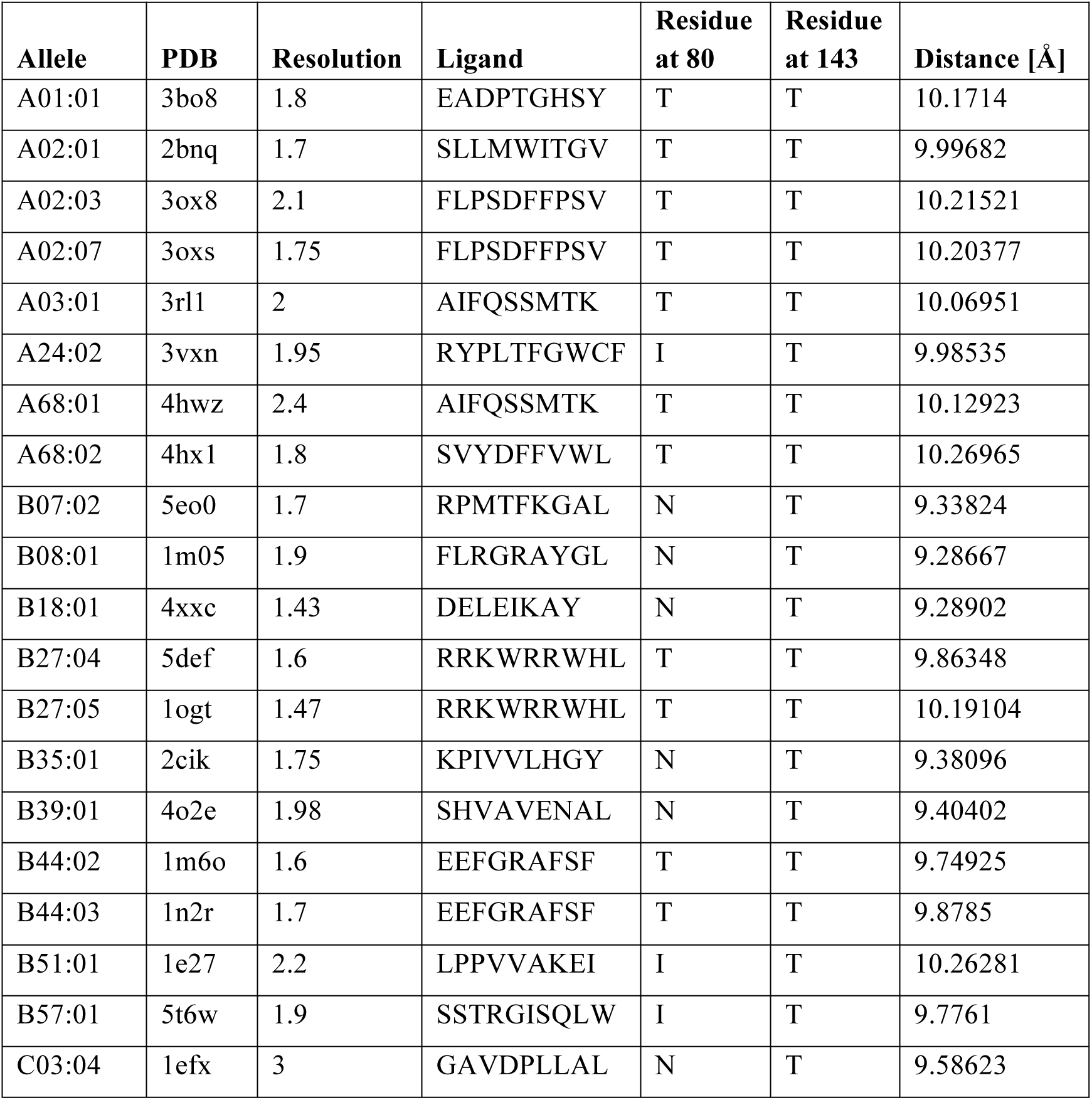
List of X-ray structures available for alleles present in our dataset of deconvoluted HLA peptidomics studies. The last column shows the distance between C-alpha atoms of residues 80 and 143 (residue numbering following A02:01 2BNQ X-ray structure).

**Table S5:** C-terminal extension predictions in *T. gondii* HLA-A02:01 ligands from (10).

|  | Number of peptides | C-terminal extension | Expected | SD | Z-score |
| --- | --- | --- | --- | --- | --- |
| 10-mers | 22 | 2 | 0.13 | 0.3 | 6.231 |
| 11-mers | 25 | 5 | 0.09 | 0.23 | 21.308 |
| 12-mers | 23 | 5 | 0.05 | 0.14 | 34.668 |

**Table S6:** Sensitivity of the C-terminal extension predictions with respect to different choices for the thresholds T_1_. Only alleles that passed the thresholds are shown.

| $T_1=1.5$ | Sample | Allele | 10-mers | C-terminal Extension | Expected | SD | Z-score |
| --- | --- | --- | --- | --- | --- | --- | --- |
| Abelin | A0203 | A0203 | 446 | 13 | 4.54 | 2.09 | 4.056 |
| Abelin | A0207 | A0207 | 424 | 17 | 5.06 | 2.12 | 5.629 |
| Ritz | HEK293 | A0301 | 1221 | 63 | 4.7 | 2.08 | 28.094 |
| Melanoma | Mel_8 | A0301 | 1283 | 25 | 13.21 | 5.04 | 2.34 |
| Abelin | A0301 | A0301 | 381 | 51 | 8.05 | 2.41 | 17.811 |
| Melanoma | Mel_15 | A0301 | 4865 | 48 | 13.27 | 4.68 | 7.414 |
| Abelin | A3101 | A3101 | 193 | 26 | 0.4 | 0.55 | 46.695 |
| Melanoma | Mel_15 | A6801 | 1944 | 38 | 14.18 | 4.74 | 5.026 |
| Lausanne | RA957 | A6801 | 4865 | 137 | 24.19 | 7.2 | 15.676 |
| Abelin | A6802 | A6802 | 357 | 11 | 4.36 | 1.68 | 3.957 |
| Abelin | B5401 | B5401 | 194 | 12 | 6.58 | 2.39 | 2.27 |

| $T_1=2.5$ | Sample | Allele | 10-mers | C-terminal Extension | Expected | SD | Z-score |
| --- | --- | --- | --- | --- | --- | --- | --- |
| Abelin | A0203 | A0203 | 446 | 10 | 2.31 | 1.42 | 5.416 |
| Abelin | A0207 | A0207 | 424 | 17 | 2.83 | 1.66 | 8.529 |
| Ritz | HEK293 | A0301 | 1221 | 62 | 0.09 | 0.29 | 212.804 |
| Lausanne | Apher1 | A0301 | 1273 | 14 | 2.5 | 1.65 | 6.952 |
| Melanoma | Mel_8 | A0301 | 1283 | 24 | 2.83 | 2.23 | 9.483 |
| Mommen | Bcell | A0301 | 2521 | 26 | 5.49 | 2.95 | 6.965 |
| Abelin | A0301 | A0301 | 381 | 45 | 1.93 | 1.39 | 31.095 |
| MCP | Fibroblast | A0301 | 859 | 17 | 2.25 | 1.89 | 7.797 |
| Abelin | A3101 | A3101 | 193 | 25 | 0.08 | 0.23 | 106.889 |
| Melanoma | Mel_15 | A6801 | 4865 | 136 | 7.65 | 3.25 | 39.49 |
| Lausanne | RA957 | A6801 | 1944 | 38 | 1.66 | 1.83 | 19.849 |
| Abelin | A6802 | A6802 | 357 | 10 | 1.57 | 0.96 | 8.736 |
| Mommen | Bcell | B0702 | 2521 | 18 | 10.17 | 2.93 | 2.676 |
| Mommen | Bcell | B2705 | 2521 | 17 | 6.22 | 2.12 | 5.093 |
| Melanoma | Mel_15 | B2705 | 4865 | 37 | 12.5 | 3.77 | 6.501 |
| Abelin | B5401 | B5401 | 194 | 12 | 4.56 | 1.94 | 3.842 |

| $T_1=\text{top 2\%}$ | Sample | Allele | 10-mers | C-terminal Extension | Expected | SD | Z-score |
| --- | --- | --- | --- | --- | --- | --- | --- |
| Abelin | A0203 | A0203 | 446 | 11 | 2.8 | 1.46 | 5.597 |
| Abelin | A0207 | A0207 | 424 | 17 | 3.64 | 1.9 | 7.037 |
| Ritz | HEK293 | A0301 | 1221 | 63 | 0.33 | 0.59 | 107.027 |
| Lausanne | Apher1 | A0301 | 1273 | 15 | 5.19 | 2.4 | 4.082 |
| Melanoma | Mel_8 | A0301 | 1283 | 25 | 6.73 | 3.74 | 4.891 |
| Mommen | Bcell | A0301 | 2521 | 27 | 11.91 | 4.23 | 3.563 |
| Abelin | A0301 | A0301 | 381 | 48 | 3.28 | 1.85 | 24.142 |
| Melanoma | Mel_15 | A0301 | 4865 | 48 | 7.8 | 3.82 | 10.535 |
| MCP | Fibroblast | A0301 | 859 | 17 | 4.69 | 2.76 | 4.464 |
| Abelin | A3101 | A3101 | 193 | 25 | 0.15 | 0.35 | 70.09 |
| Lausanne | RA957 | A6801 | 1944 | 38 | 7.06 | 3.12 | 9.932 |
| Melanoma | Mel_15 | A6801 | 4865 | 137 | 16.55 | 5.7 | 21.137 |
| Abelin | A6802 | A6802 | 357 | 11 | 1.98 | 1.15 | 7.837 |
| Abelin | B5401 | B5401 | 194 | 12 | 5.55 | 1.96 | 3.284 |

**Table S7:** Crystallographic data collection and refinement statistics (* Values in parentheses correspond to the highest resolution shell).

| **Data Collection** |  |
| --- | --- |
| PDB ID | **6EI2** |
| Protein/Ligand | HLA-A*68:01/β2M/peptide |
| Space group | P2<sub>1</sub>2<sub>1</sub>2<sub>1</sub> |
| Cell dimensions: a, b, c (Å) | 59.06 80.25 110.88 |
| $\alpha$, $\beta$, $\gamma$ (deg) | 90.00 90.00 90.00 |
| Resolution* (Å) | 1.61 (1.70-1.61) |
| Unique observations* | 68846 (9873) |
| Completeness* (%) | 99.9 (99.5) |
| Redundancy* | 6.6 (6.6) |
| Rmerge* | 0.094 (0.602) |
| I/ σI* | 11.3 (2.8) |
| **Refinement** |  |
| Resolution (Å) | 1.61 |
| R<sub>work</sub> / R<sub>free</sub> (%) | 15.9 / 18.1 |
| Number of atoms<br>(protein/other/water) | 3165 / 30 / 442 |
| B-factors (Å<sup>2</sup>)<br>(protein/other/water)21.85 | 19.95 / 33.69 / 30.45 |
| r.m.s.d bonds (Å) | 0.016 |
| r.m.s.d angles (°) | 1.714 |
| Ramachadran Favoured (%) | 97.33 |
| Allowed (%) | 2.67 |
| Disallowed (%) | 0.00 |

